# Age-related variability in network engagement during music listening

**DOI:** 10.1101/2023.03.24.534123

**Authors:** S. Faber, A.G. Belden, P. Loui, A.R. McIntosh

## Abstract

Listening to music is an enjoyable behaviour that engages multiple networks of brain regions. As such, the act of music listening may offer a way to interrogate network activity, and to examine the reconfigurations of brain networks that have been observed in healthy aging. The present study is an exploratory examination of brain network dynamics during music listening in healthy older and younger adults. Network measures were extracted and analyzed together with behavioural data using a combination of hidden Markov modelling and partial least squares. We found age- and preference-related differences in fMRI data collected during music listening in healthy younger and older adults. Both age groups showed higher occupancy (the proportion of time a network was active) in a temporal-mesolimbic network while listening to self-selected music. Activity in this network was strongly positively correlated with liking and familiarity ratings in younger adults, but less so in older adults. Additionally, older adults showed a higher degree of correlation between liking and familiarity ratings consistent with past behavioural work on age-related dedifferentiation. We conclude that, while older adults do show network and behaviour patterns consistent with dedifferentiation, activity in the temporal-mesolimbic network is relatively robust to dedifferentiation. These findings may help explain how music listening remains meaningful and rewarding in old age.

## Background

Brain function changes with age across multiple spatial scales. The brain can be thought of as a series of overlapping functional networks where each network is a collection of brain regions that act in concert over time. With age, regions that were once nodes in densely-connected functional networks may become disconnected while regions in previously distinct functional networks may become more connected (Grady et al., 2016), though whether this reconfiguration of functional network boundaries is adaptive or maladaptive remains unclear. In healthy older adults, networks that were once well-defined and responded preferentially to a particular stimulus or set of conditions begin to activate (or to fail to deactivate) less discerningly in a process known as dedifferentiation (Grady et al., 2012; Rieck et al., 2017).

In music listening, there is behavioural evidence of age-related perceptual changes that may serve as a behavioural counterpart to the dedifferentiation seen in network brain dynamics. Music is reported as more broadly pleasant with age (a positivity effect, Bones & Plack, 2015; Groarke & Hogan, 2019; Laukka & Juslin, 2007; Lima & Castro, 2011), and perceptual features also become less distinct with age, with higher correlations observed between perceived arousal and valence in older adults (Vieillard et al., 2012). This blurring of the lines between the perceived pleasantness and dimensions of a musical signal might indicate underlying network changes, but do not seem to affect the music listening experience negatively.

Musical sounds are complex stimuli that, using building blocks of timbre, tone, pitch, rhythm, melody, and harmony, can engender expectancy and surprise to make us laugh, cry, dance, sing, and reminisce. As musical stimuli are complex and hierarchically organized, brain responses to music are likewise complex and hierarchical, with many temporally-dependent overlapping processes. Features extracted from musical signals stimulate activity in multiple brain regions (Alluri et al., 2012; Burunat et al., 2017; Williams et al., 2022), and networks, including the default mode network (DMN; Wilkins et al., 2014; Koelsch et al., 2022; Taruffi et al., 2017) and reward networks (Fasano et al., 2022).

Multivariate statistical modelling tools provide us with a unique opportunity to observe and describe whole-brain network activity in a data-driven way. Working in network space, where the smallest unit of measurement is a network, allows us to examine the shifting patterns of brain activity that accompany music, which has the potential to add nuance that cannot be seen when looking at isolated regions of interest. This approach may also be of value in understanding the neural foundation of age-related perceptual changes, and may shed light on why music is so salient in clinical populations (Cuddy & Duffin, 2005; Leggieri et al., 2017; Särkämö et al., 2014; Thaut et al., 2020, Matziorinis & Koelsch, 2022).

Where older adults show network reconfigurations compared to younger cohorts in rest and during cognitive tasks, what can music reveal about the aging brain? In the present exploratory study, we studied age differences in network-level dynamics during familiar and novel music listening in a cohort of healthy younger and older adults. We aim to demonstrate age-related changes in network dynamics using a novel analysis paradigm comprising hidden Markov modelling and partial least squares analyses.

## Methods

Networks were estimated using hidden Markov modelling (HMM) and analyses were completed using partial least squares (PLS). We chose HMM rather than a seed-based or canonical network analysis (see Bressler & Menon, 2010) in an effort to base our analyses on data-driven patterns as much as possible. A substantial advantage of HMM is that it derives networks from patterns in the original data without the constraints of canonical network boundaries or specified time windows.

A brief outline of data collection is included here. For a detailed description of participant recruitment, study protocol, and data acquisition, please see Quinci et al. (2022) and Belden et al. (2023).

### Participants

Participants were right-handed, cognitively healthy younger (N = 44, 11 males, mean age = 19.24, SD = 1.92) and older (N = 27, 13 males, mean age = 67.34, SD = 8.27) adults with normal hearing established via audiogram. Inclusion criteria included normal hearing, successful completion of MRI screening, and a minimum age of 18 for younger adults and 50 for older adults. Exclusion criteria comprised medication changes 6 weeks prior to screening, a history of any medical condition that could impair cognition, a history of chemotherapy in the preceding 10 years, or any medical condition requiring medical treatment within three months of screening. Data from two younger participants were excluded following data collection due to problems with the ratings apparatus. Ethics approval was granted by the Northeastern University Institutional Review Board and all research was conducted consistent with the Declaration of Helsinki.

### Procedure

Prior to data collection, participants completed a screening call with researchers to confirm their eligibility for the study, and to collect a list of six songs that are familiar and well-liked by the participant. Following screening, eligible participants completed a battery of neuropsychological tests, structural and functional MRI scans, and a blood draw. The present study focuses on the fMRI data; other aspects of the results are in preparation and will be described in separate reports.

### Data acquisition

All scans took place at Northeastern University. Functional scans were acquired with a Siemens Magnetom 3T scanner with a 64-channel head coil. The total scan time for task data was 11.4 minutes with continuous acquisition at a fast TR of 475 ms over 1440 volumes. A resting state scan was also performed with these parameters, and findings will be reported in a future manuscript. T1 images were captured, but will not be discussed in detail in this manuscript.

Task fMRI consisted of a block of resting state followed by music presentation (24 excerpts, each played for 20 seconds). Musical excerpts were either familiar and well-liked self-selected music (6/24), or experimenter-selected music chosen to be popular or possibly recognizable (10/24), or novel including excerpts purpose-composed for research purposes (8/24). Stimuli were presented randomly and following each 20 second musical excerpt, participants were asked to rate their familiarity and liking of the excerpt for two seconds each, using 4-point Likert scales.

### Data pre-processing

Functional MRI data were pre-processed using the TVB-UKBB pipeline detailed by Frazier-Logue et al. (2022). T1 images were registered to the Montreal Neurological Institute T1 template. Functional data pre-processing was done using a pipeline using the FMRIB Software Library (FSL; Woolrich et al., 2009), including the fMRI Expert Analysis Tool (FEAT, version 6.0). Within the pipeline, pre-processing of functional data comprised gradient echo fieldmap distortion correction, motion correction using MCFLIRT, and independent component analysis (ICA) artifact classification using MELODIC and FIX.

We assembled an ICA training set for non-cerebral artifact detection. ICA reports from 16 participants per age group were visually inspected for noisy vs. clean components and manually annotated. Subsequent participants’ ICA reports were cleaned using this training set. The processed datasets were down-sampled to 220 regions of interest using the Schaefer-Tian 220 parcellation, which provides ample spatial resolution of auditory regions and subcortical structures (Schaefer et al., 2017, Tian et al., 2020). Regional time series data were normalized to control for between-subject amplitude differences and exported to MatLab (MathWorks, 2019) for Hidden Markov Model estimation and analysis.

### Network Estimation

To estimate networks, we used the HMM-MAR Toolbox (Vidaurre et al., 2017, 2018). The estimation uses ROI time series data and calculates the K networks that best describe the entire dataset. It then allocates each time window to the single best-fitting network within the original time series. HMM, as a dimensionality reduction technique, returns states (hereafter referred to as networks) that can be used to observe how networks interact over time.

The output from HMM is a time series showing the most prominent network at each timepoint. From this timeseries, it is possible to calculate fractional occupancy and state-wise transitional probability (Vidaurre et al., 2017). Fractional occupancy is the proportion of the total number of timepoints each network was occupied during a time series task, and shows a particular network’s prominence during target time windows. Transitional probability shows the most likely patterns of steps from one network to another. Thus, both are related measures, but contain different information about how the networks interact.

We estimated HMMs with variable K values between 3 and 20. We found the estimations with 4 and 7 states to provide the most optimal model-derived free energy values (see Vidaurre et al., 2017; Vidaurre et al., 2018). Partial least squares analyses showed statistically significant effects for both estimations with comparable effect sizes (see Fasano et al., 2022). We further interrogated the spatial properties of the states in each estimation by computing a dot product of the normalized state means, finding that the spatial properties of the states in the estimation with 7 states were well-represented in the estimation with 4 states. We ultimately chose the 4 state estimation as it provided a single state with activity in temporal and mesolimbic regions together. Temporal and mesolimbic region activity has been previously related to auditory reward (Salimpoor et al., 2011, Fasano et al., 2020), including prior analyses of subsets of the present data (Belden et al., 2023, Quinci et al., 2022).

The K networks identified by the HMM estimation are shown in Figure 1 (cortical regions only) and the regions of interest are detailed in Table 1. Where this analysis did not use canonical network-based seeds, we assigned anatomical labels to the networks based on the taxonomy of functional brain networks consistent with the wider network literature (Uddin et al, 2019). The functional properties of these states will be addressed in the discussion.

**Figure 1:**
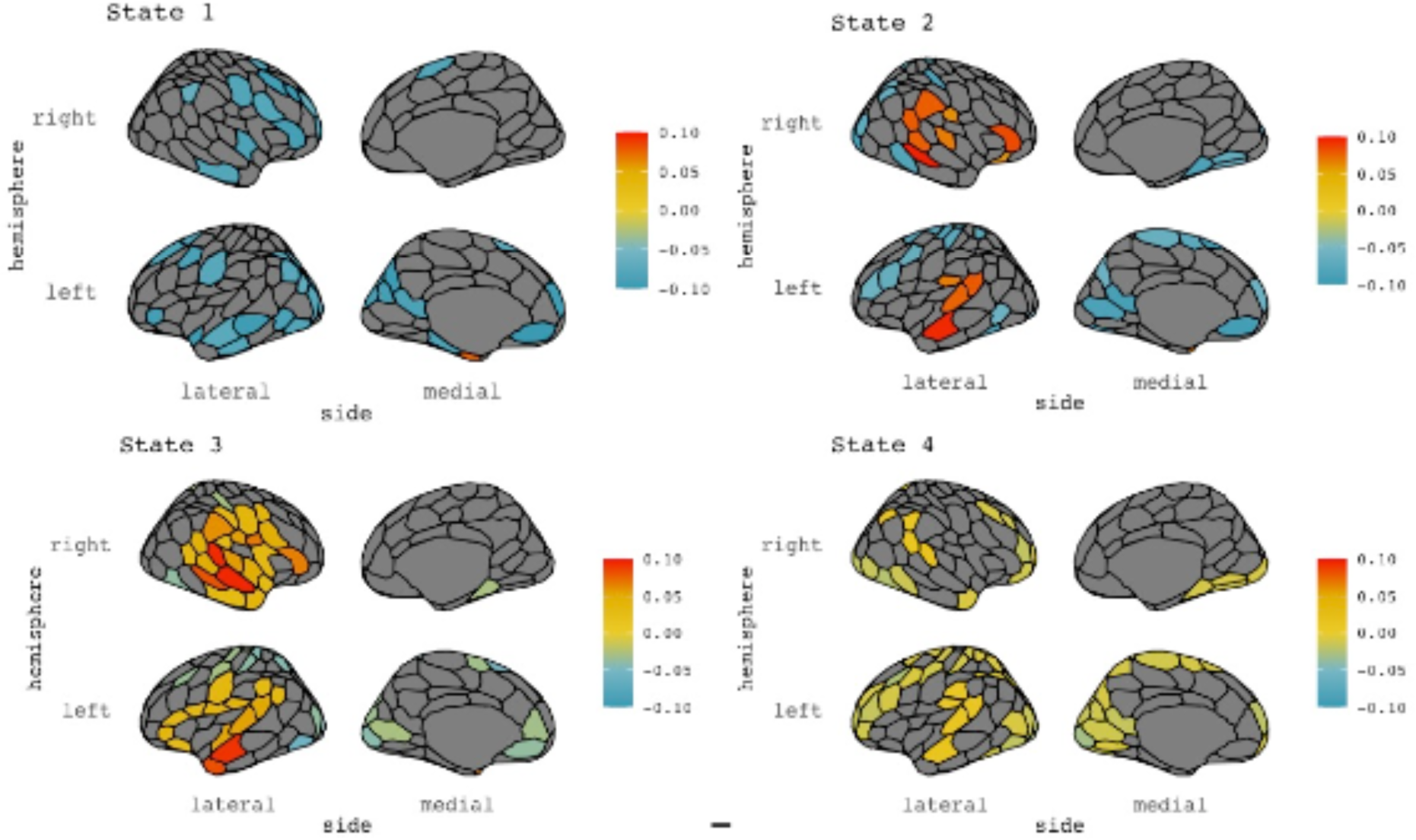
Mean activity plots returned from HMM analysis. Colours represent relative activity of the states and all have been normalized within-state. See Table 1 for subcortical regions not displayed here.

**Table 1:** Regions of interest and network labels from HMM analysis. Network labels are based on the work of Uddin et al. (2019).

| State | Main Regions | Network |
| --- | --- | --- |
| 1 | Bilateral middle-frontal and left temporal regions.<br>Subcortical regions include the bilateral temporal pole, left nucleus accumbens, and right hippocampal body | Medial frontoparietal network |
| 2 | Bilateral temporal and frontal regions | Temporal network |
| 3 | Bilateral temporal and mesolimbic regions<br>Subcortical regions include the left globus pallidus, left hippocampal body, right putamen, and right hippocampal tail | Temporal mesolimbic network |
| 4 | Bilateral superior frontal and middle parietal regions | Frontoparietal network |

### PLS

We used partial least squares (PLS) to analyze between- and within-group differences on the HMM-extracted measures. PLS is a multivariate analysis technique that uses singular value decomposition to quantify the relationship(s) between data matrices and experimental features, in this case, fractional occupancy and transitional probability measures. In these analyses, we used mean-centred PLS to analyze group and task differences using the HMM-extracted measures and the within-subject relation of the measure to participant liking and familiarity ratings. To emphasize group main effects, we performed mean-centred analyses subtracting the overall grand mean from the group means. To focus on task main effects and task by group interactions, secondary mean centred analyses were performed, subtracting the group mean from the task mean within each group (i.e., rendering the group main effect zero).

PLS analysis returns mutually-orthogonal latent variables (LVs) that describe group and/or task effects. Each LV’s statistical significance and reliability are calculated via permutation testing and bootstrap estimation, respectively with a statistical threshold of *p* <.05. The reliability and strength of the group or task effects is depicted through the confidence interval estimation of LV scores for all participants, where LV scores are the dot-product of subject data and LV weights. LV weights themselves are evaluated for reliability through bootstrap ratios of the weight divided by its estimated standard error, which can be interpreted as a z-score for the corresponding confidence interval (see McIntosh & Lobaugh, 2004).

## Results

Prior to HMM decomposition, we tested for sex differences using mean-centred PLS on each participant’s average functional connectivity matrix from the music listening task. No significant sex-related differences were found. Following these analyses, we ran additional PLS analyses to test for sex effects in fractional occupancy and transitional probability, returning no significant effects. Data were subsequently pooled together for the remainder of the analysis.

### Fractional Occupancy

We extracted average fractional occupancy for each participant, and fractional occupancy for each participant for each category of musical excerpt (self-selected, experimenter-selected popular, and experimenter-selected novel) and used PLS to observe differences in fractional occupancy across age groups and stimuli categories. Mean-centred PLS analysis returned one significant LV (*p* = .024) showing an age effect, with younger adults showing higher fractional occupancy in the temporal network (network/state 2), and older adults showing higher fractional occupancy in the frontoparietal network (network/state 4).

When divided into stimulus categories and analyzed for task main effects and task-by-group interactions, mean-centred PLS analysis returned one significant LV (*p* < .01, Figure 2) showing an effect of self-selected music vs experimenter-selected music on fractional occupancy in the temporal-mesolimbic network (network 3). Fractional occupancy is higher for this network while listening to self-selected music (music that is highly familiar and well-liked) in both younger and older adults. Fractional occupancy for the temporal network (network 2) is higher when listening to experimenter-selected music. Both effects are qualitatively more reliable in younger adults based on confidence intervals (Figure 3A).

**Figure 2:**
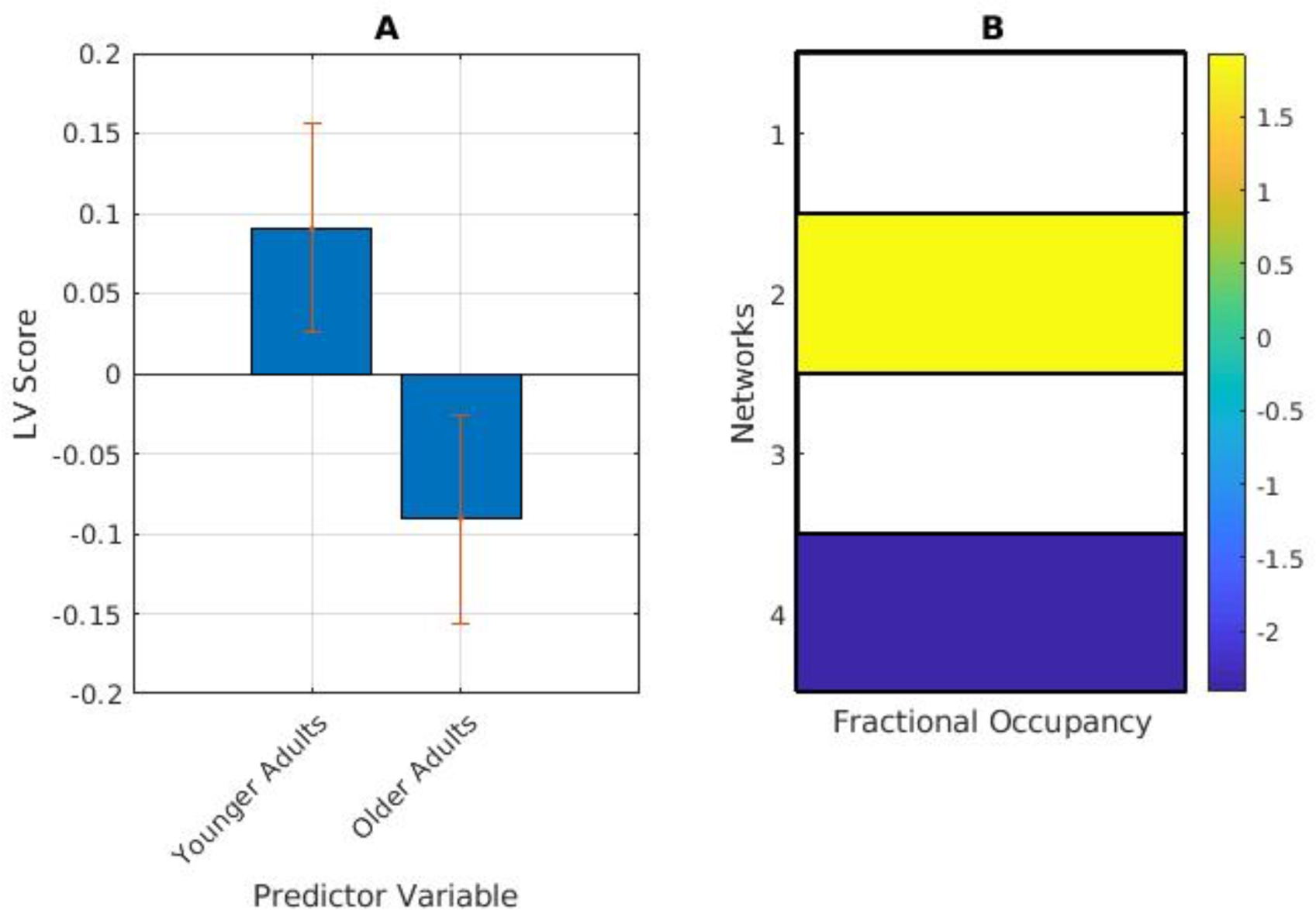
Age-related differences in fractional occupancy (FO). (A) PLS contrasts between age groups in music listening. Error bars were calculated using bootstrap resampling and reflect the 95% confidence interval. The contrasts show an age effect on FO (B), with the higher FO in network 2 in younger adults, and higher FO in network 4 in older adults. The colour scale represents the bootstrap ratio for each network.

**Figure 3:**
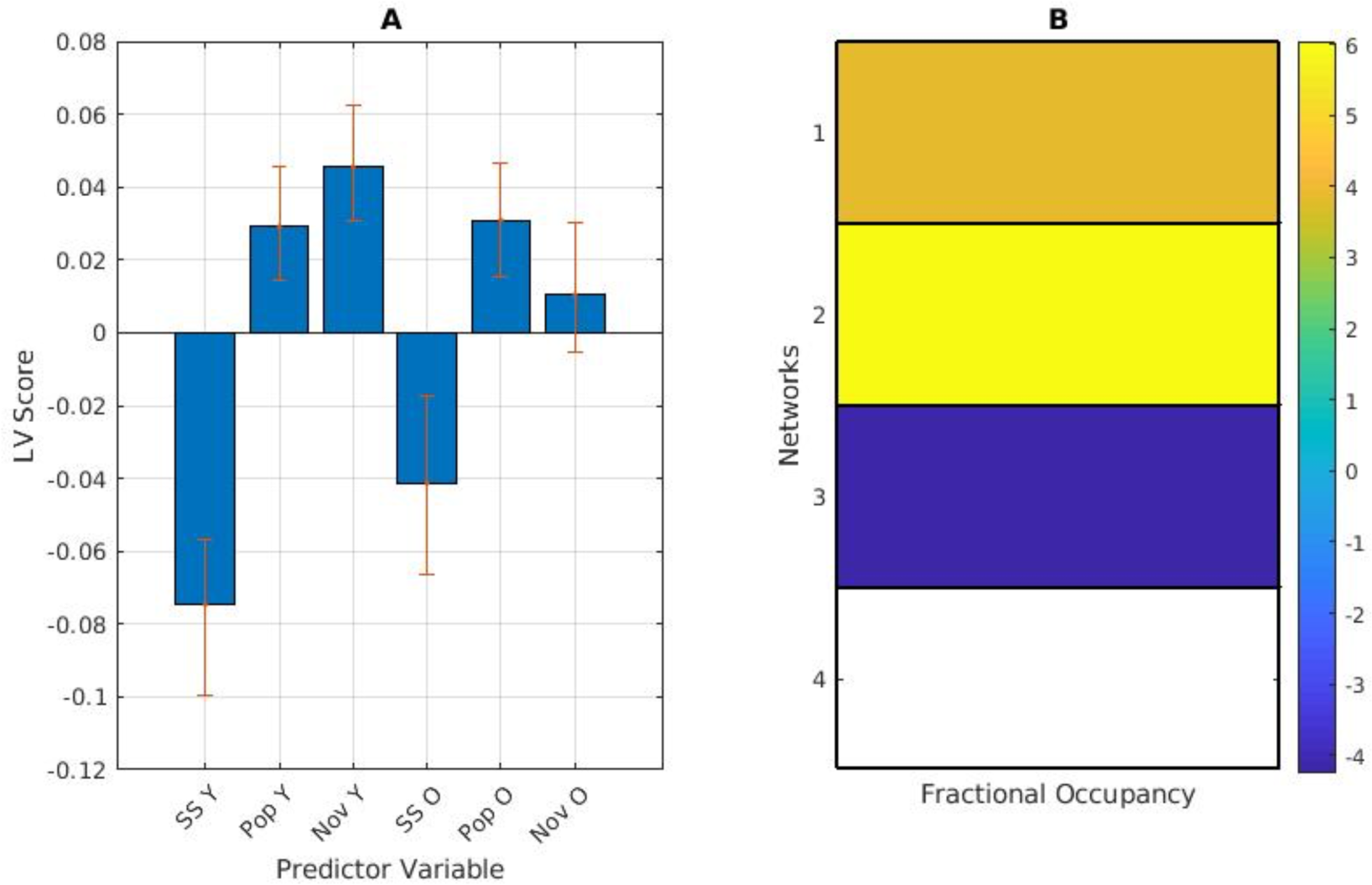
(A) PLS contrasts between age groups in stimuli category and fractional occupancy (FO). Error bars were calculated using bootstrap resampling and reflect the 95% confidence interval. The contrasts show a stimulus-type effect on FO in both age groups (B), with the higher FO in network 3 in both groups during self-selected music listening (SS Y and SS O), and higher FO in network 2 during experimenter-selected music listening (Pop and Nov delineating popular and novel excerpts respectively).

### Transitional Probability

We next examined the transitional probability matrices for differences in network interaction on average and between the different stimulus categories. Important to note: the data being analyzed is the directional likelihood of transitioning from each network to each other network. Rather than looking at networks by themselves, these results show the link or edge that connects each network to each other network.

The averaged transitional probability mean-centred PLS returned one significant LV (*p* < .001), showing a contrast between younger and older adults, with younger adults more likely than older adults to transition into the temporal network (network 2) from other networks, and less likely than older adults to transition to the frontoparietal network (network 4) from the temporal network (network 2) In examining network persistence (the likelihood of staying in a network), younger adults were more likely to stay in the temporal network when listening to experimenter-selected music (Figure 4).

**Figure 4:**
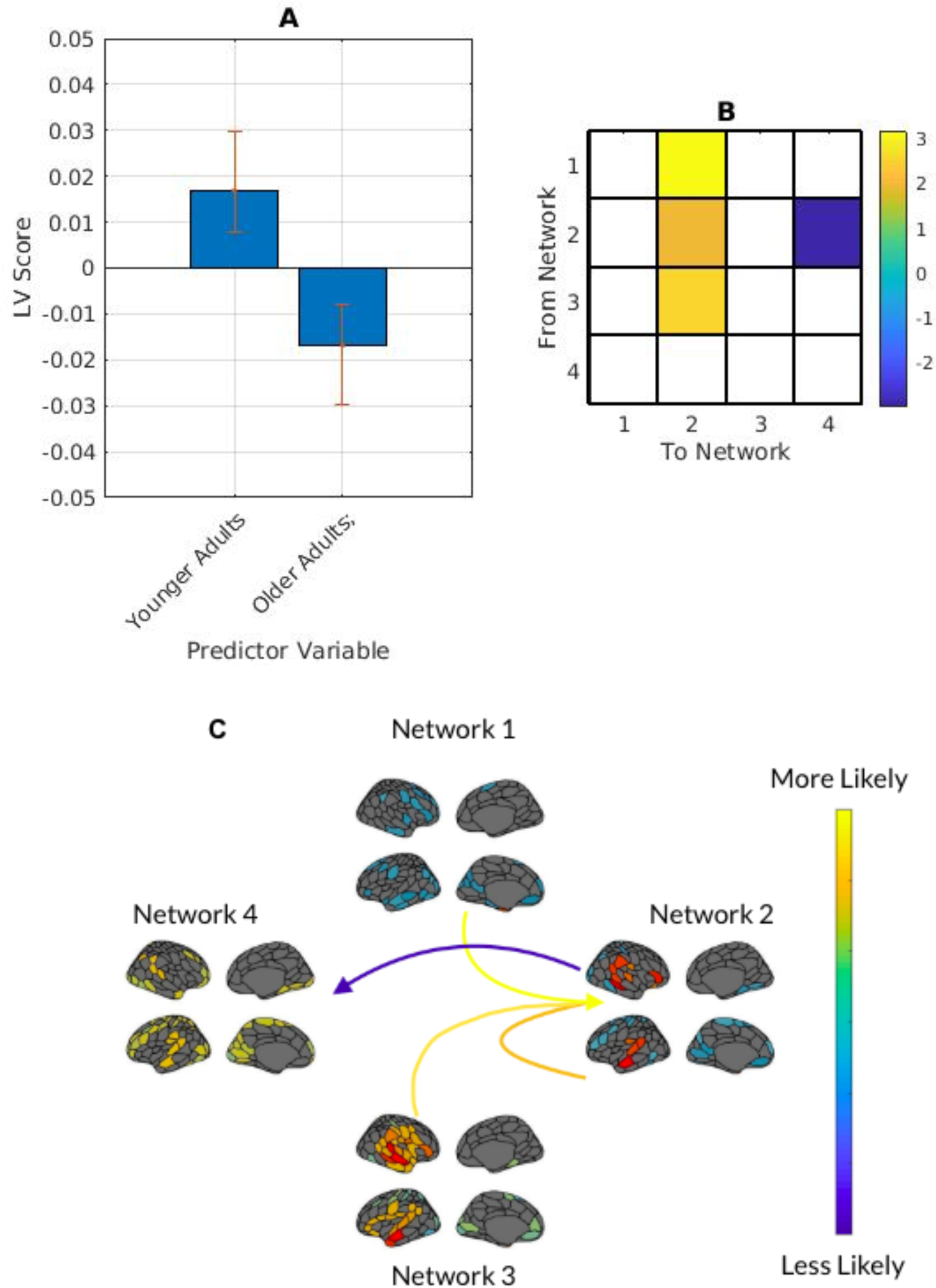
(A) PLS contrasts between age groups and transitional probability (TP). Error bars were calculated using bootstrap resampling and reflect the 95% confidence interval. The contrasts show an age effect on TP in both age groups (B), with younger adults more likely to transition into network 2 from networks 1, 2, and 3 than older adults; and less likely to transition to network 4 from network 2 than older adults (C). The colour scale represents the bootstrap ratio for each network.

When divided into stimulus categories and analyzed for task main effects and task-by-group interactions, both groups were more likely to transition from the temporal network to the temporal-mesolimbic and frontoparietal networks during self-selected music listening. In experimenter-selected music, both groups were most likely to transition from the temporal-mesolimbic network to the temporal network, but this effect was more pronounced in younger adults. In examining network persistence (the likelihood of staying in a network), all participants were more likely to stay in the temporal-mesolimbic network when listening to self-selected music and more likely to stay in the temporal network when listening to experimenter-selected music. When analyzed within age, older participants did not show a significant network persistence pattern in the temporal network during experimenter-selected music (Figure 5).

**Figure 5:**
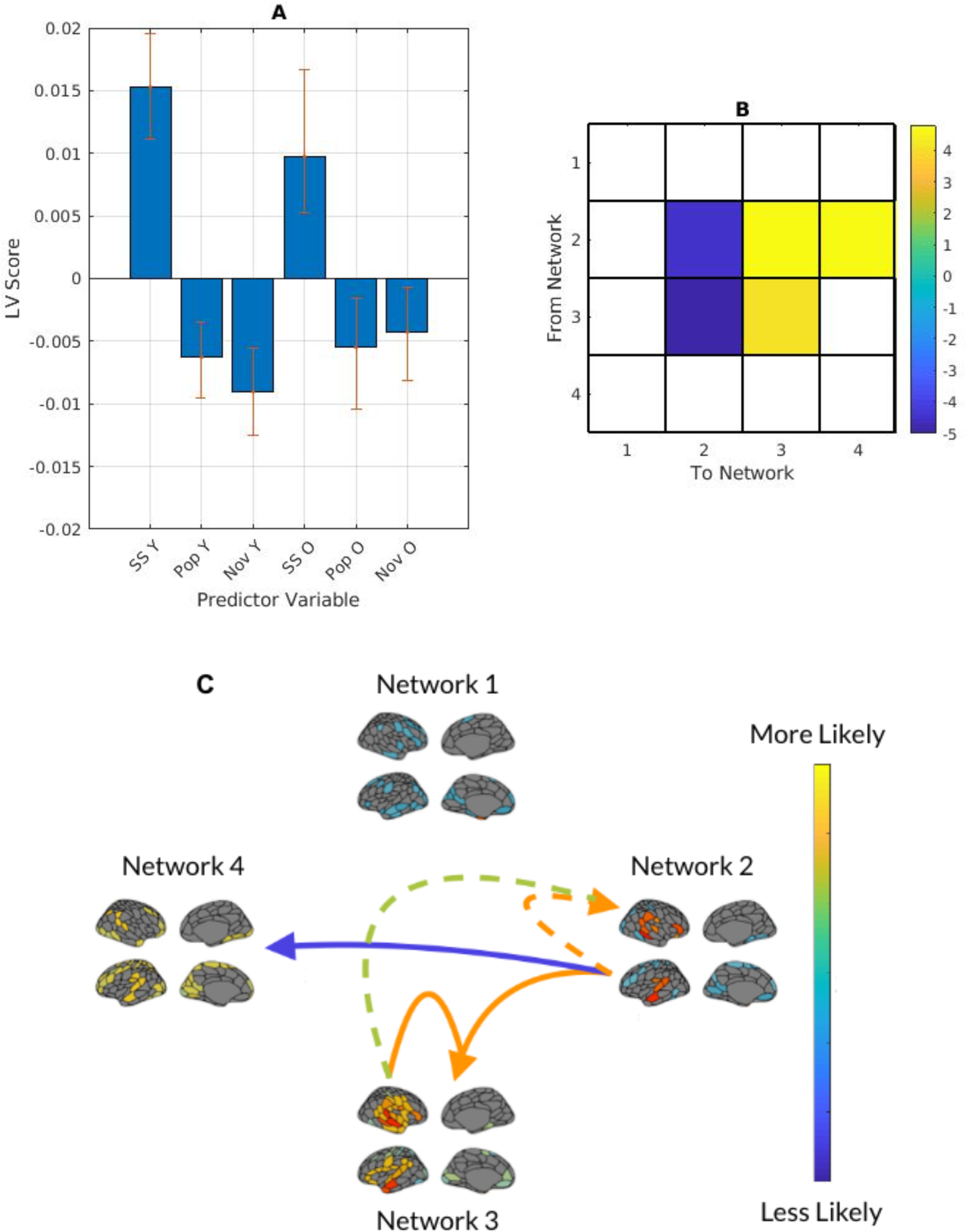
(A) PLS contrasts between age groups in stimulus category and transitional probabilities. SS refers to self-selected music, Pop and Nov refers to popular and novel experimenter-selected music. Error bars were calculated using bootstrap resampling and reflect the 95% confidence interval. The contrasts (B) show a stimulus-type effect on transitional probability (TP), illustrated with the TP magnitude in panel C. Panel C shows the between-network TP with solid lines representing self-selected music and dashed lines representing experimenter-selected music.

**Figure 6:**
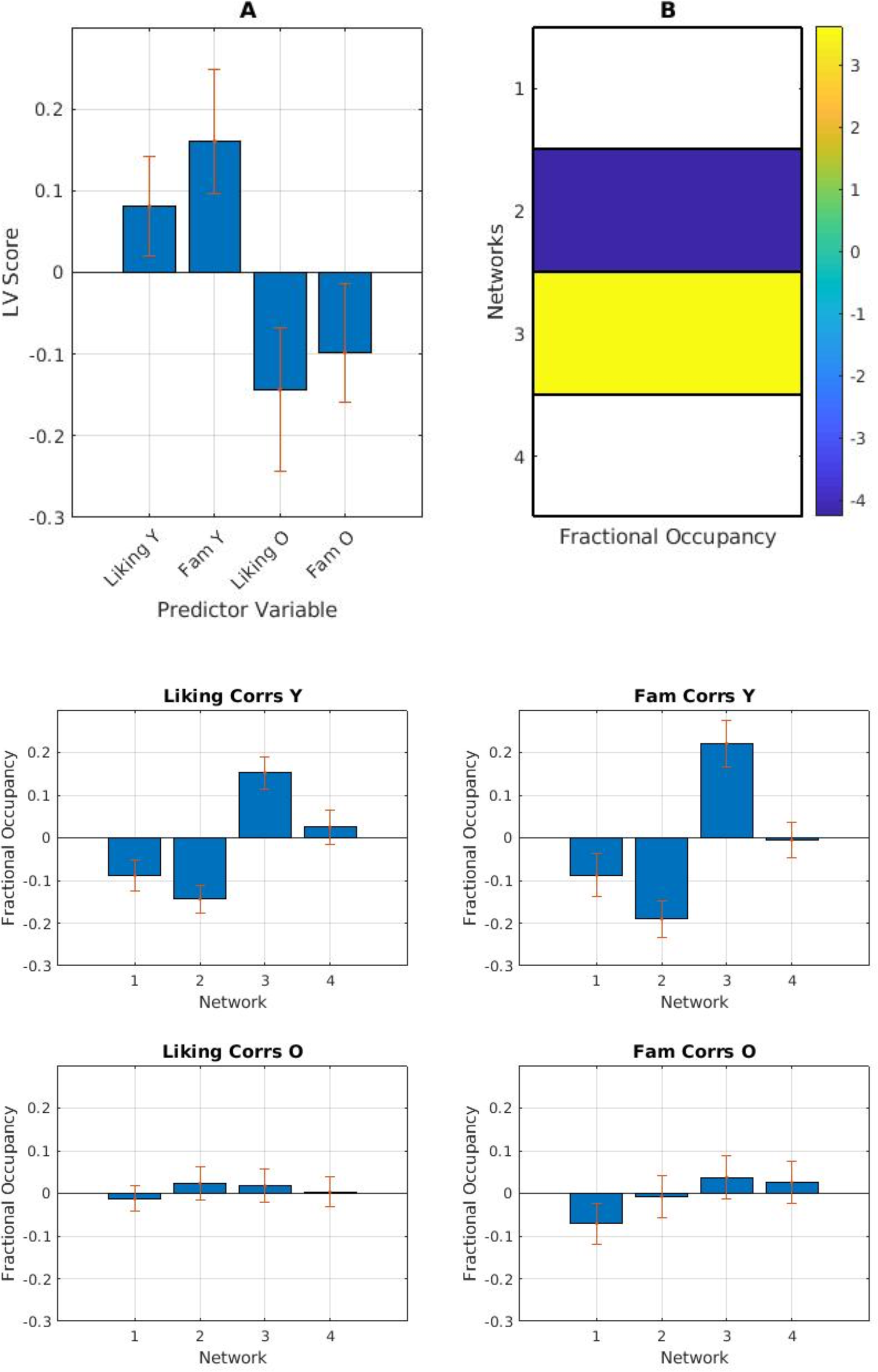
(A) PLS contrasts between age groups in stimulus category and fractional occupancy. Error bars were calculated using bootstrap resampling and reflect the 95% confidence interval. The contrasts show an age effect on correlations between liking and familiarity (Fam) and network fractional occupancy (B), illustrated with the relevant magnitude in panel C.

### Effects of liking and familiarity on brain measures

We next analyzed the network fractional occupancy and transitional probability matrices with participants’ liking and familiarity ratings. We correlated liking and familiarity ratings for each excerpt with fractional occupancy for each participant. Initial mean-centred PLS analysis returned no significant LVs. Following this analysis, we ran the PLS centred to the overall grand mean to allow for a full factorial analysis: group main effect, task main effect and group-by-task interactions.

The results from the full factorial PLS returned one significant LV (*p* < .001) showing the contrast between age groups. In younger adults, the temporal-mesolimbic network featured prominently, showing a greater positive correlation than other networks with both liking and familiarity. Older adults showed a more ambiguous correlation between liking and familiarity and fractional occupancy in the temporal network (Figure 7).

**Figure 7:**
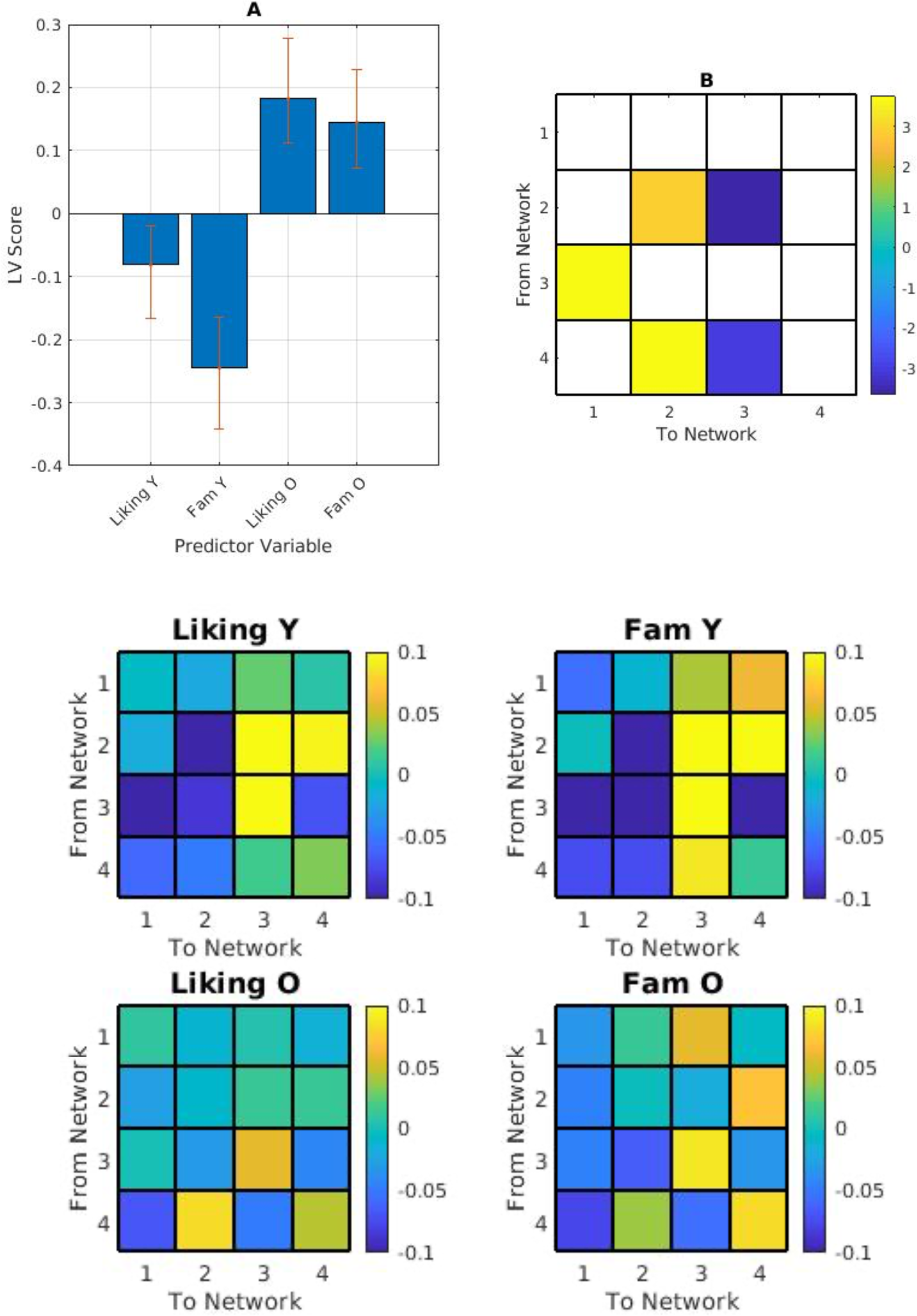
(A) PLS contrasts between age groups in stimulus category and transitional probabilities. Error bars were calculated using bootstrap resampling and reflect the 95% confidence interval. The contrasts (B) show an age effect on correlations between liking and familiarity (Fam) and network transitional probability, illustrated with the relevant magnitude in panel C. The colour scale represents the bootstrap ratio for each network.

We next vectorized the excerpt-wise transitional probability matrices for each participant, and correlated them with each participant’s piece-wise liking and familiarity ratings, returning two transitional probability-correlation matrices per participant: liking*transitional probability and familiarity*transitional probability.

A full factorial PLS consistent with the above analysis returned one significant LV (*p* < 0.001) showing an age effect. Younger adults’ liking and familiarity ratings were more strongly positively correlated with the likelihood of transitioning to the temporal-mesolimbic network from the temporal and frontoparietal networks. Younger adults’ ratings were more strongly negatively correlated with persistence in the temporal-mesolimbic network, and the likelihood of transitioning from the temporal-mesolimbic network to the medial frontoparietal network. Transitioning from the frontoparietal network to the temporal network was more positively correlated with ratings in older adults, and more negatively correlated with ratings in younger adults (Figure 7).

Within-age mean-centred PLS analyses did not return any significant LVs.

### Liking and familiarity behavioural ratings

Finally, we examined the ratings themselves. Mean-centred PLS showed older adults rated excerpts as significantly less familiar than younger adults (*p* < .01). However, they did not significantly differ in liking ratings. Mean-centred PLS also showed older adults’ liking and familiarity data were significantly more highly correlated than younger adults (*r* = 0.57 for older adults and *r* = 0.43 for younger adults, PLS *p* < .01).

## Discussion

Music listening engages multiple brain networks that may reorganize in multiple ways as we age. While there are well-documented effects of music listening on auditory and reward networks and auditory-motor networks, less is known about how music listening may encourage persistence within networks, or transitions between networks. Treating data-driven brain networks as units of analysis, we detailed age-related similarities and differences in network occupancy and between-network transitional probabilities during music listening. The two most commonly-featured networks in these analyses were the temporal and temporal-mesolimbic networks. Activity in temporal-mesolimbic regions overlaps with auditory-reward network activity (see Wang et al., 2020), while temporal regions are firmly affiliated with auditory processing (Belfi & Loui, 2019).

Both younger and older adults showed the highest fractional occupancy in the temporal-mesolimbic network while listening to self-selected music compared to experimenter-selected music. These stimuli were selected by participants to be familiar and well-liked, and auditory-reward network activation for preferred music has been well-documented in prior studies (Salimpoor et al., 2011, Fasano et al., 2020), including on a subset of these data (Quinci et al., 2022). This network was active for experimenter-selected music as well, though to a lesser extent than self-selected music, particularly in younger adults.

When looking at the transitional probability matrices, self-selected music was again linked to persistence in the temporal-mesolimbic network and a greater probability of transition to this network from the temporal network in both age groups. Experimenter-selected music was linked to higher persistence in the temporal network and a greater probability of transition to the temporal network from the temporal-mesolimbic network in both age groups, indicating that music listening employs a distributed network of frontal and temporal regions; but to engage mesolimbic structures, a degree of liking and familiarity is needed.

However, when analyzed separately, group differences were more obvious. Older subjects showed an increased likelihood of persistence in the temporal network during experimenter-selected music, but this effect was less reliable than in younger adults. Older adults also showed an increased likelihood of transitioning to the temporal-mesolimbic network from the medial frontoparietal network in self-selected music. This network shares many regions with the default mode network (DMN; Uddin et al., 2019). The DMN is implicated in listening to liked (Wilkins et al., 2014; Pereira et al., 2011) and timbrally rich music (Alluri et al., 2012), and is less attenuated during cognitive tasks with age (Rieck et al., 2017). One possible explanation is that older adults are less likely to transition from the medial frontoparietal network to the temporal network during music listening than younger adults, instead remaining in the medial frontoparietal network until transitioning to the temporal-mesolimbic network while a younger adult may transition from the medial frontotemporal network to the temporal network.

The older adult transitional probability matrices showed more transitions to the temporal-mesolimbic network during experimenter-selected music, which could indicate an age-related shift in between-network dynamics. Former pathways (in this case, the likelihood of transitioning from an auditory reward network to an auditory perception network during unfamiliar music, or staying in an auditory perception network during unfamiliar music) reconfigure in favour of consistency across multiple types of music involving the temporal mesolimbic network. This is consistent with earlier findings that network functional specificity declines in favour of a more standard set of responses to multiple stimuli types (Rieck et al., 2020).

In younger adults, liking and familiarity ratings were correlated with fractional occupancy in the temporal and temporal mesolimbic networks, with the temporal network most occupied when familiarity and liking are low and the temporal mesolimbic network most occupied when familiarity and liking are high. In older adults, correlations between fractional occupancy and liking and familiarity ratings are more ambiguous, indicating a reconfiguration of network engagement related to aging. Correlations between ratings and transitional probabilities were consistent with this pattern: younger adults’ likelihood of transitioning into the temporal and temporal-mesolimbic networks were more strongly correlated with liking and familiarity than older adults who showed a more diffuse pattern.

Older adults showed high fractional occupancy in the temporal-mesolimbic network during all music types. This difference could be because older adults show less differentiation between liking and familiarity during novel music listening. If familiarity is lower among older adults, but liking is consistent with younger adults, it is possible that older adults would engage a different network response to music that is unfamiliar but liked. Liking and familiarity are more positively correlated in older adults than younger adults, consistent with earlier findings on age-related blunting of emotional intensity and liking (where stimuli are consistently rated as less extremely pleasant and unpleasant. See Baird et al., 2020; Groarke & Hogan, 2019; Laukka & Juslin, 2007).

While these results offer a promising look into capturing age-related changes in network-level dynamics in naturalistic behaviours, there are several areas for further inquiry. To more fully examine age, future studies could include a more continuous range of participants, particularly those in middle adulthood to disambiguate age and cohort effects. While this study did not focus on music and memory, future work could include a measure of music-related memory (see Jakubowski & Eerola, 2022) to disambiguate group differences due to memory and lived experience. The methods presented here were in effort to identify networks most relevant to this dataset in a data-driven way. This approach, while advantageous in presenting nuanced fluctuations in network membership, may prove challenging to reconcile with the canonical network literature. Future work could employ both canonical and data-driven methods to directly examine network membership and behaviour in an effort to link both methodological approaches.

These observations could illustrate the broader pattern of the network dynamics of music listening, and the age-related reorganization of these networks. For older adults, the temporal network becomes less finely tuned to liking and familiarity, while the temporal mesolimbic network remains active. There are several exciting implications of these findings. The first is in studying naturalistic behaviours in “network space”: investigating the behaviours and interactions of networks as behaviour unfolds. The need to understand the brain as a complex, dynamic system, one that is continually adapting to its surroundings, has been the topic of much discussion (see McIntosh & Jirsa, 2019; Calhoun et al., 2014). The brain is more than a collection of regions and its emergent properties can be captured in fascinating detail using music. Though the methods presented here are not unique to music, we also hope to present music as a viable stimulus to interrogate higher cognition.

In the same way that the brain is not merely a collection of regions, music is more than a simple collection of notes. It is ubiquitous in the human experience (Savage, 2019; Cross & Morley, 2010) but has yet to experience its renaissance in cognitive neuroscience. There are good reasons for this: music data contain many layers of information from the content of the signal itself to the content of the memories or the quality of movement it generates in the listener. However, the scientific potential of music is too beguiling to ignore. Here is a stimulus that, unlike rest, has a rich, externally-measurable temporal structure that, unlike traditional task paradigms, does not require extensive training or fortitude to endure. It combines the best of both worlds with the added benefit of being accessible to clinical populations in ways that other tasks, especially those reliant on language, are not.

By examining music’s network properties, we present a data-driven methodological framework for future hypothesis-driven studies of musical behaviour while offering an alternative to traditional paradigms that is externally measurable, ecologically valid, and accessible to those with cognitive decline or who are non-verbal.

